# Asymmetric post-translational modifications regulate the intracellular distribution of unstimulated STAT3 dimers

**DOI:** 10.1101/546523

**Authors:** Beatriz Cardoso, Ricardo Letra-Vilela, Catarina Silva-Almeida, Ana Maia Rocha, Fernanda Murtinheira, Carmen Rodriguez, Vanesa Martin, Federico Herrera

**Affiliations:** Cell Structure and Dynamics Laboratory, Instituto de Tecnologia Quimica e Biologica (ITQB-NOVA), Universidade Nova de Lisboa, Oeiras, Portugal; Instituto Universitario de Oncología del Principado de Asturias (IUOPA) and Departamento de Morfología y Biología Celular, Facultad de Medicina, c/Julian Claveria, 33006 Oviedo, University of Oviedo, Spain

**Keywords:** STAT3, bimolecular fluorescence complementation, dimerization, post-translational modifications, acetylation

## Abstract

Signal Transducer and Activator of Transcription 3 (STAT3) is a ubiquitous and pleiotropic transcription factor that plays essential roles in normal development, immunity, response to tissue damage and cancer. We have developed a Venus-STAT3 bimolecular fluorescence complementation (BiFC) assay that allows the visualization and study of STAT3 dimerization and protein-protein interactions in living cells. Inactivating mutations on residues susceptible to post-translational modifications (K49R, K140R, K685R, Y705F and S727A) changed significantly the intracellular distribution of unstimulated STAT3 dimers when the dimers were formed by STAT3 molecules that carried different mutations. Our results indicate that asymmetric post-translational modifications on STAT3 dimers could constitute a new level of regulation of STAT3 signaling.

**Significance:** The Signal Transducer and Activator of Transcription 3 (STAT3) transcription factor plays key roles in development, immunity, cancer or response to stress or damage. All previous studies on STAT3 dimerization work on a homogenous pool of STAT3 molecules, where all STAT3 molecules are equally modified (i.e.“symmetric”). However, this is highly unlikely in a complex intracellular environment, as post-translational modifications do not necessarily occur with complete efficiency or simultaneously. We demonstrate that asymmetric post-translational modifications change the intracellular distribution of STAT3 dimers more strikingly than symmetric ones. This could mean a new level of regulation of STAT3 activity, and therefore a new possible therapeutic target. Our results could be highly relevant for other protein complexes regulated by post-translational modifications beyond STAT3.

## Introduction

The Signal transducer and activator of transcription 3 (STAT3) is a conserved transcription factor that plays key roles in development, immunity, response to injury and cancer (1,2). STAT3 homodimerization, post-translational modification (PTM) and intracellular location are key events in its biological functions. STAT3 is canonically activated by phosphorylation at Y705 upon stimulation with a variety of cytokines and growth factors. However, unstimulated STAT3 also dimerizes, is found in the nucleus, binds to DNA and controls the transcription of a specific set of genes, different from phosphorylated STAT3 (1–5). STAT3 can be also found in the mitochondria, where it is necessary for normal activity of the electron transport chain (6,7). This function is independent of its activity as a transcription factor and Y705 phosphorylation, but dependent on S727 phosphorylation (1,6,7). Other PTMs can regulate the behavior and function of STAT3, such as acetylation at K49 or K685 (3,8,9) or dimethylation at K49 or K140 (10,11). Although dimethylation of the K49 or K140 residues is induced by stimulation with cytokines and is favored by STAT3 phosphorylation, there is basal K49 (but not K140) dimethylation in unstimulated STAT3 (10), and the same happens with STAT3 acetylation (8,9). Literature on STAT3 generally assumes that STAT3 homodimers are formed by two identically modified molecules. However, this is highly unlikely in a complex intracellular context, as PTMs do not occur in all the pool of STAT3 molecules at the same time or with the same efficiency. Here, we aimed at determining the relative contribution of residues K49, K140, K685, Y705 and S727 to the dimerization and intracellular distribution of STAT3 homodimers.

## Material and Methods

We developed a suit of plasmids to study STAT3 homodimerization in living cells, based on bimolecular fluorescence complementation (BiFC)(12). Briefly, the cDNA sequence of STAT3-alpha was fused to the sequence of two complementary, non-fluorescent fragments of the Venus protein (Venus 1, amino acids 1-157; and Venus 2, amino acids 158-238)(Fig. 1A), and inserted in a pcDNA 3.3 TOPO backbone (Invitrogene). When STAT3 dimerizes, the Venus fragments are brought together and reconstitute the fluorophore, the fluorescence being proportional to the amount of dimers (Fig. 1A). Deletion mutants lacking the C-terminus (DelCT) were produced by PCR-mediated subcloning using full-length Venus-STAT3 BiFC constructs as templates. The original lysine (K) residues on positions 49, 140 and 685 were replaced by arginine (R) residues, the tyrosine (Y) residue on position 705 by phenylalanine (F) and the serine (S) residue on position 727 by alanine (A)(Fig. 1A). Supplementary Table I shows the primers used for cloning and mutagenesis were. Detailed methods for site-directed mutagenesis, cell culture and transfection, fluorescence microscopy, flow cytometry and immunoblotting were described elsewhere (12). All BiFC constructs were deposited in Addgene (https://www.addgene.org/). HeLa human cervix adenocarcinoma cells and HEK293 human embryonic kidney cells were acquired from ATCC (references CRM-CCL-2 and CRL-1573, respectively), Leukemia inhibitory factor (LIF) from R&D systems (MN, USA), and Stattic from Selleckchem (TX, USA).

**Figure 1.**
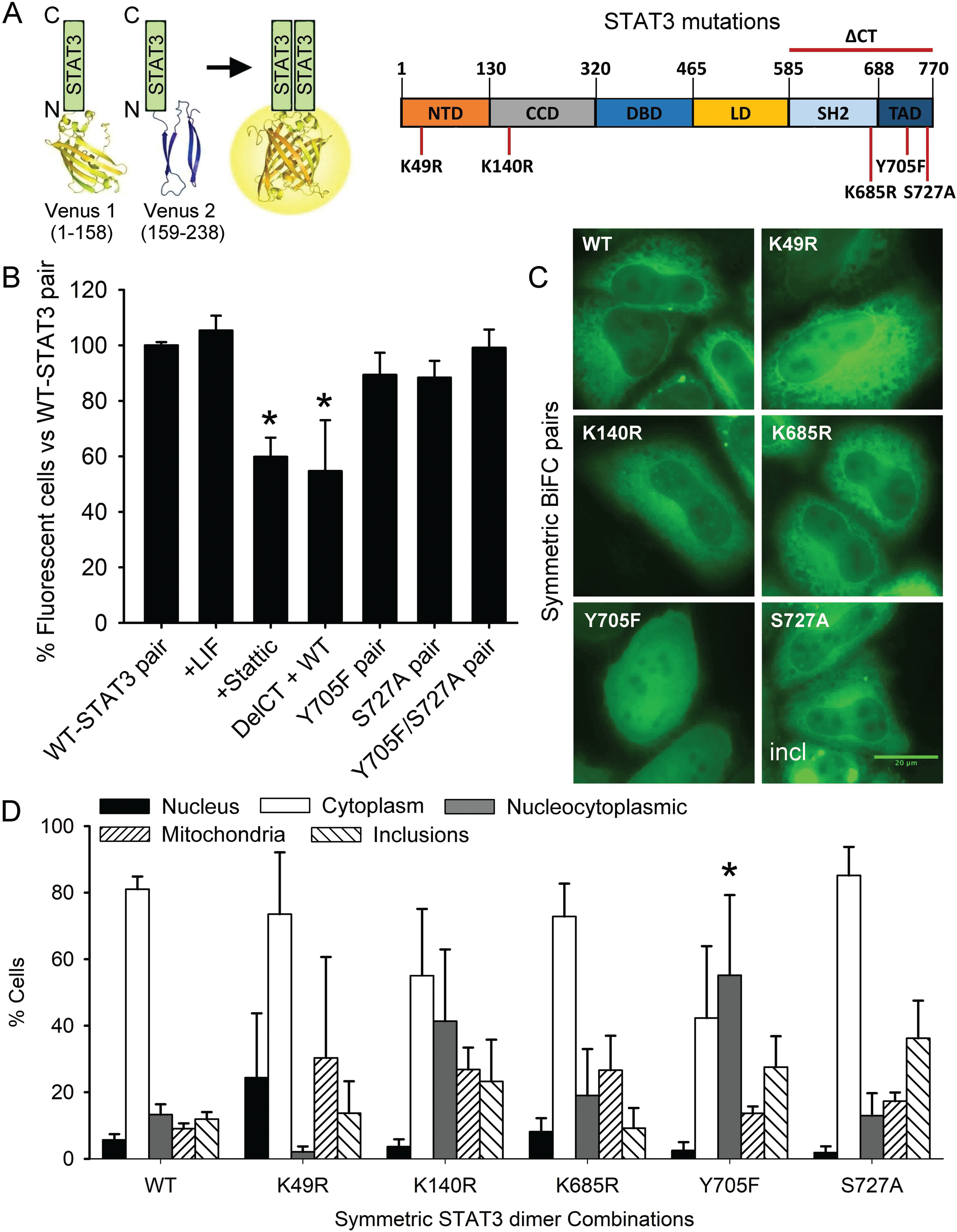
A Venus-STAT3 BiFC system allows the visualization and study of STAT3 homodimers in living cells. A, Venus BiFC fragments constituted by amino acids 1-158 (Venus 1, V1) and 159-238 (Venus 2, V2) were fused to the N-terminus of the STAT3 sequence in two independent constructs. K49, K140, K685, Y705 and S727 residues can be post-translationally modified, and were inactivated in both V1- and V2-STAT3 constructs by site-directed mutagenesis. B, Wild-type Venus-STAT3 constructs produced fluorescence in HeLa cells, and it was monitored by flow cytometry 24 hours after transfection with BiFC constructs. Incubation with Leukemia Inhibitory Factor (100 ng/ml) for 2 hours in the absence of serum or the presence of the indicated drugs or mutant BiFC pairs (n=3, p<0.05). Results were normalized versus the Wild-type STAT3 pair (100%). C, Microscopy pictures of representative cell phenotypes in the different symmetric combinations of BiFC Venus-STAT3 constructs (Incl, inclusions; Scale bar, 20μm). D, Percentage of cells displaying fluorescence predominantly in the Nucleus (black bar), predominantly in the Cytosol (white bar), homogeneously distributed in cytoplasm and nucleus (nucleocytoplasmic, grey bar), in the mitochondria or in non-mitochondrial inclusions. Data are shown as the average ± SEM of n=12 (Wild-type, WT) or n=3 (rest of combinations) independent experiments. Statistical analysis was carried out by means of a One-way ANOVA followed by a Bonferroni test adjusted for multiple comparisons. *, sign. vs the symmetric Wild-type STAT3 pair, p<0.05.

## Results

Transfection of HEK293 or HeLa cells with the wild-type (WT) pair of Venus-STAT3 BiFC constructs led to successful expression of the chimeric proteins V1-STAT3 and V2-STAT3 (Figs. 1B and 1C, Suppl. Fig. 1). Fluorescence was primarily cytoplasmic in both cell lines, with low but visible nuclear signal (Fig. 1C, Suppl. Fig. 1B). Incubation with Leukemia Inhibitory Factor (LIF, 100 ng/ml) induced STAT3 phosphorylation and translocation to the nucleus in HEK293 and HeLa cells (Suppl. Fig. 1B-C), but it did not enhance STAT3 dimerization (Fig. 1B, Suppl. Fig. 1D). Incubation with STAT3 inhibitor Stattic (5 μM) or removal of the C-terminus containing the SH2 domain partially prevented STAT3 dimerization (Fig. 1B), consistent with previous reports (13,14). On the other hand, single or double Y705F/S727A phosphoresistant mutants did not decrease fluorescence (Fig. 1B). These results support relevant evidence indicating that STAT3 dimerization is actually independent of phosphorylation (1,5,14). The behavior of the Venus-STAT3 BiFC system is therefore consistent with previous reports for tagged STAT3.

In order to analyze the intracellular location of unstimulated STAT3 homodimers, we classified cells qualitatively in three categories that are mutually exclusive (their sum is 100% of cells), according to the relative intensity and location of the fluorescence signal (Fig. 1C-1D): 1) predominantly in the cytoplasm (e.g. WT pair), 2) predominantly in the nucleus (e.g. upon LIF induction, Suppl. Fig. 1B), or 3) homogeneously distributed through nucleus and cytoplasm (e.g. Y705F pair). We also determined the percentage of cells with mitochondrial signal or intracellular inclusions (Suppl. Fig. 2). Although changes in patterns of STAT3 dimer distribution were observed in several symmetric BiFC pairs, only Y705F and S727A pairs induced significant increases in the percentage of cells with homogeneous nucleocytoplasmic fluorescence or inclusions, respectively (Fig. 1D).

We made use of the unique properties of our BiFC system to determine the relative contribution of each residue to the dimerization and intracellular distribution of unstimulated STAT3 dimers. We combined all possible inactivating PTM mutations with each other, but no combination had a consistent effect on STAT3 dimerization as determined by flow cytometry (Suppl. Fig. 3). However, the intracellular distribution of STAT3 homodimers was significantly altered by specific combinations of STAT3 molecules (Fig. 2A). Unlike the K49R symmetric pair, K49R asymmetric combinations dominantly induced an increase in cells with homogeneous nucleocytoplasmic fluorescence at the expense of cytoplasmic location (Fig. 2A), similar to the Y705F symmetric pair. K140R- or K685R-containing pairs showed some tendency to shift cytoplasmic location towards nucleus, but only the K140R+S727A combination achieved significance. This phenotype was almost identical to the Y705F+S727A asymmetric pair (Fig. 2A).

**Figure 2.**
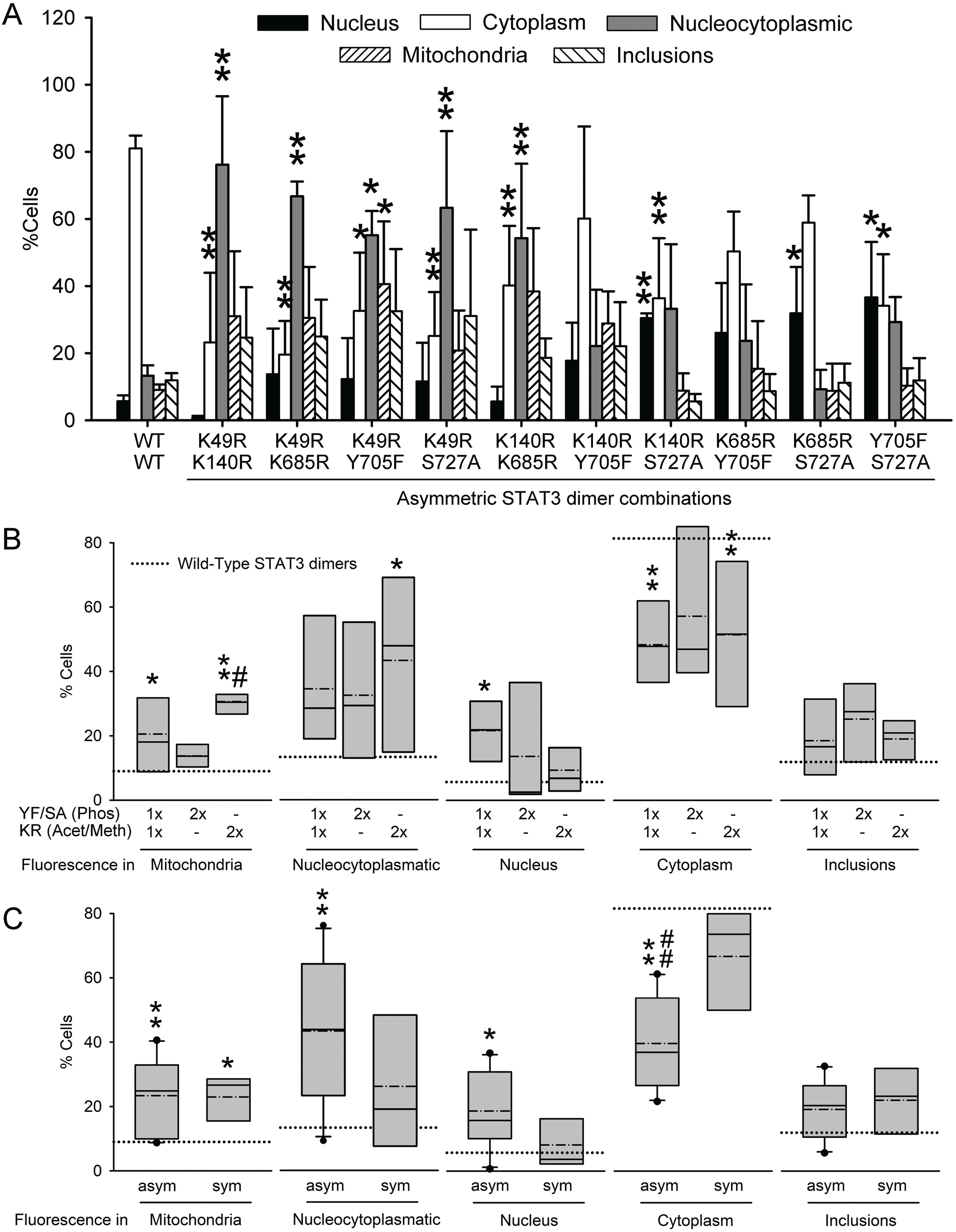
Asymmetric STAT3 PTMs regulate intracellular distribution of STAT3 homodimers. A, Intracellular distribution of fluorescence in asymmetric combinations of Venus-STAT3 BiFC constructs (and the WT symmetric pair as reference). Data are shown as the average of n=12 (Wild-type, WT) or n=3 (rest of combinations) independent experiments ± SEM. Statistical analysis was carried out by means of a One-way ANOVA followed by a Bonferroni test adjusted for multiple comparisons. *, significant vs the symmetric Wild-type STAT3 pair, *, p<0.05, ** p<0.01. B-C, The same original data, but pooled according to the number and nature of substitutions (B) or the symmetry or asymmetry of substitutions (C) in the STAT3 homodimer, and represented as box plots. The limits of the boxes represent the smallest and largest values, the straight line represents the median, the dashed line represents the average, and the dotted line represents the average for wild-type STAT3 pair. Statistical analysis was carried out on the Average ± SEM of each pool of data (1xYF/SA:1xKR, n=6; 2xYF/SA, n=3; 2xKR, n=6; sym, n=5; asym, n=10). Statistical analysis was carried out by means of a One-way ANOVA followed by a Bonferroni test adjusted for multiple comparisons. *, significant vs the symmetric Wild-type STAT3 pair, *, p<0.05, ** p<0.01; #, significant vs 2xYF/SA substitution (in B) or the symmetric mutant pairs (in C), #, p<0.05, ##, p<0.01.

We then pooled and analyzed all results according to the number and type of PTM mutations present in the each BiFC pair. Combinations carrying any one (asymmetric) or two K-R substitutions (symmetric or asymmetric) significantly increased mitochondrial translocation, while decreasing the percentage of cells with STAT3 dimers predominantly in the cytoplasm (Fig. 2B). Asymmetric combinations of one K-R substitution and one phosphoresistant mutant also increased nuclear translocation, but only 2xK-R combinations increased homogeneous nucleocytoplasmic distribution. Combinations carrying any two phosphoresistant mutations (symmetric or asymmetric) had no significant effect on cellular distribution of STAT3 homodimers (Fig. 2B). These results indicate that specific asymmetric PTMs on STAT3 dimers can prevent their nuclear import/export. This was later confirmed by pooling the data according to whether the STAT3 pair was symmetric or asymmetric in their PTM mutations (Fig. 2C). We found that only asymmetric PTM mutant combinations increased nucleocytoplasmic or nuclear distribution at the expense of decreasing cytoplasmic localization of STAT3 homodimers. Asymmetric combinations were also sufficient to produce an increase in mitochondrial localization of STAT3 dimers (Fig. 1C).

## Discussion

Our results indicate that asymmetric PTMs could constitute a new level of regulation of unstimulated STAT3 behavior and function. Most STAT3 molecules are not phosphorylated in the absence of extracellular stimuli, and this proportion is reversed shortly after cytokines bind to their corresponding membrane receptors (Suppl. Fig. 1C). However, cells often show small amounts of phosphorylated STAT3 in resting state (Suppl. Fig. 1C) and, conversely, cytokine-stimulated STAT3 induces the *de novo* transcription of new STAT3 molecules that are not necessarily phosphorylated (1,2). This indicates that unphosphorylated and phosphorylated STAT3 should coexist at similar levels in many situations, and the literature presents evidence that this could be equally true for other STAT3 PTMs induced by cytokines (8–10).

To the best of our knowledge, there is no direct empirical evidence in the literature showing that asymmetric STAT3 dimers actually happen in living cells, and such demonstration would be currently extremely difficult from a technical point of view, even *in vitro*. Previous studies most often rely on systems that do not differentiate between monomers and dimers (2,3,6–11), and/or that produce a single population of STAT3 molecules, either mutated or normal (5,14,15). And yet, in the crowded and diverse intracellular environment, the probability for two identical STAT3 molecules to form a dimer (or for a dimer to be modified in both molecules simultaneously and in the same residues) should be low, although it could certainly be enhanced by either total absence or presence of stimuli. Our results point in this exciting direction, and open a series of interesting questions. If asymmetric STAT3 dimers actually happen, do they regulate specific sets of genes? Do they enable gradation of STAT3 transcriptional or mitochondrial activities? And if they do not happen, how do cells manage to achieve perfectly symmetrical STAT3 dimers with such high efficiency? Given the essential roles of STAT3 in development, immunity, tissue stress and cancer, addressing these questions could have important implications for the diagnosis, treatment and understanding of a wide spectrum of human pathologies.

## Acknowledgements

The authors thank the Advanced Imaging Unit from Gulbenkian Science Institute and Dr. Sixto Herrera for support with bioimaging and flow cytometry, and statistics, respectively. FH was supported by Project LISBOA-01-0145-FEDER-007660 (Cellular Structural and Molecular Microbiology) funded by FEDER funds through COMPETE2020 - Programa Operacional Competitividade e Internacionalização (POCI) and by national funds through Fundação para a Ciência e Tecnologia (FCT, Ref. IF/00094/2013/CP1173/CT0005 and PTDC/MED-NEU/31417/2017). RLV and FM were supported by fellowships from FCT (Refs. PD/BD/128163/2016 and SFRH/BD/133220/2017, respectively).

Contribution
BC is responsible for most microscopy studies, and RV for cloning, mutagenesis and flow cytometry studies, as well as part of microscopy analyses. CSA and AMR are responsible for the first optimization experiments with the BiFC system and LIF experiments, a small part of which are shown in Suppl. Fig. 1. FM contributed to the microscopy, flow cytometry and immunoblotting at different stages of the project. CR and VM contributed to the analysis, presentation and interpretation of data. FH had the original idea, designed the experiments, arrange the final figures and analyzed the data, coordinated the project and wrote the manuscript.

## Abbreviations

BiFC: bimolecular fluorescence complementation
PTMs: post-translational modifications

