## Supplementary material for "Asymmetric post-translational modifications regulate the intracellular distribution of unstimulated STAT3 dimers"

**Supplementary Information**

**Supplementary Table I.** Primers for mutagenesis or PCR cloning.

| | | |
|---|---|---|
| STAT3 K49R | Fwd | CCCCTTGGATTGGGAGTCAAGATTG |
| | Rev | CAATCTTGACTCCCAATCCAAGGGG |
| STAT3 K140R | Fwd | GGTGACGGAGACACAGCAGATGCTG |
| | Rev | CAGCATCTGCTGTCTCTCCGTCACC |
| STAT3 K685R | Fwd | GAGGCATTCGGAAGGTATTGTCGGCC |
| | Rev | GGCCGACAATACCTTCCGAATGCCTC |
| STAT3 Y705F | Fwd | CAGGTAGCGCTGCCCCATTCCTGAAGACCAAGTTTATC |
| | Rev | GATAAACTTGGTCTTCAGGAATGGGGCAGCGCTACCTG |
| STAT3 S727A | Fwd | CATTGACCTGCCGATGGCACCCCGCACTTTAGATTC |
| | Rev | GAATCTAAAGTGCGGGGTGCCATCGGCAGGTCAATG |
| V1-STAT3 DelCT | Fwd | ACTAGCTAGCATGGTGAGCAAGGGCGAGGA |
| | Rev | CTATGGATCCTTAGTTCCAAAGGGCCAGGA |

**Supplementary Figure Legends**

**Supplementary Figure 1. The Venus-STAT3 BiFC system allows to monitor nuclear translocation of STAT3 dimers upon stimulation with LIF.** A, Flow cytometry charts showing that V1-STAT3 and V2-STAT3 BiFC constructs do not produce fluorescence by themselves, but they fluoresce when they are transfected together. B, Representative fluorescence microscopy pictures of HEK293 or HeLa cells transfected with Venus-STAT3 constructs and incubated in the presence or absence of Leukemia Inhibitory Factor (LIF, 100 ng/ml) for 2 h or 15 min, respectively. STAT3 dimers translocate to the nucleus of stimulated cells, as expected (Scale bar, 20µm). C, Immunoblotting of STAT3 in nuclear extracts from HEK293 or HeLa cells transfected with Venus-STAT3 constructs and incubated in the presence or absence of LIF (100 ng/ml) for 2 h or 15 min, respectively. Three bands are observed, corresponding to V1-STAT3 (1), V2-STAT3 (2) and endogenous STAT3 (E). D, LIF does not induce further STAT3 dimerization, as determined by flow cytometry. The histogram (right) shows that there is almost perfect overlap between the fluorescent signal emitted by cells in the presence or absence of LIF.

**Supplementary Figure 2. Examples of cells showing mitochondrial localization of STAT3 dimers versus STAT3 inclusions.** Mitochondrial localization of STAT3 dimers was monitored by co-localization of fluorescent foci with Mitotracker Red. When there were fluorescent foci that did not co-localize with Mitotracker Red we classified them as “Inclusions”.

**Supplementary Figure 3. Selected PTMs (symmetric or asymmetric) do not regulate STAT3 homodimerization in unstimulated cells.** The graph represents the percentage of fluorescent cells observed in each Venus-STAT3 BiFC combination by means of flow cytometry. The numeric code is represented in the table below the graph. Errors represent the Standard Error. Statistical analysis was carried out by means of a One-way ANOVA. No combination produced significant changes.

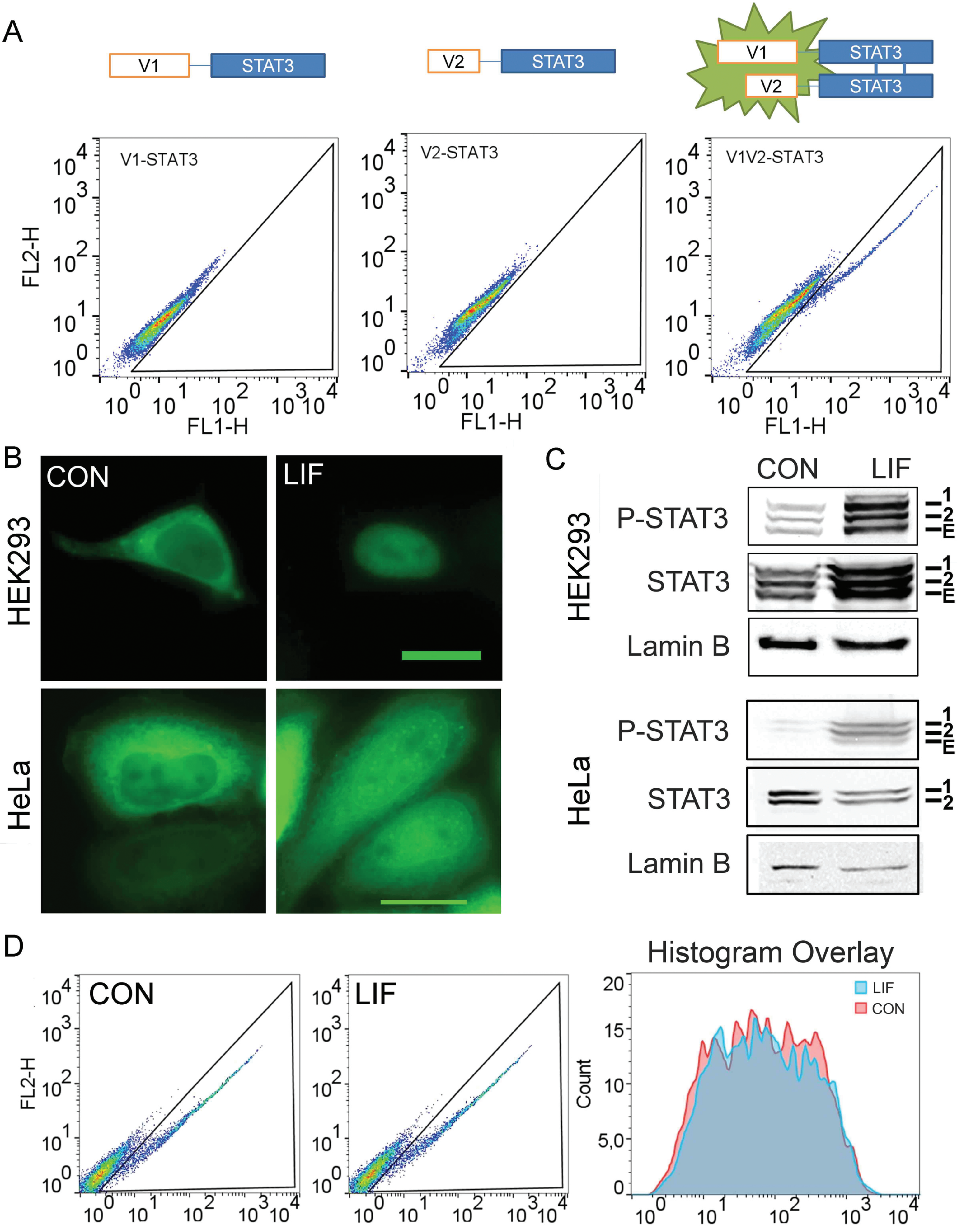
A
V1
STAT3
V2
STAT3
V1
STAT3
V2
STAT3
V1-STAT3
V2-STAT3
V1V2-STAT3
FL2-H
FL1-H
B
CON
LIF
HEK293
HeLa
C
CON
LIF
P-STAT3
STAT3
Lamin B
1
2
E
D
CON
LIF
FL2-H
Histogram Overlay
LIF
CON
Count

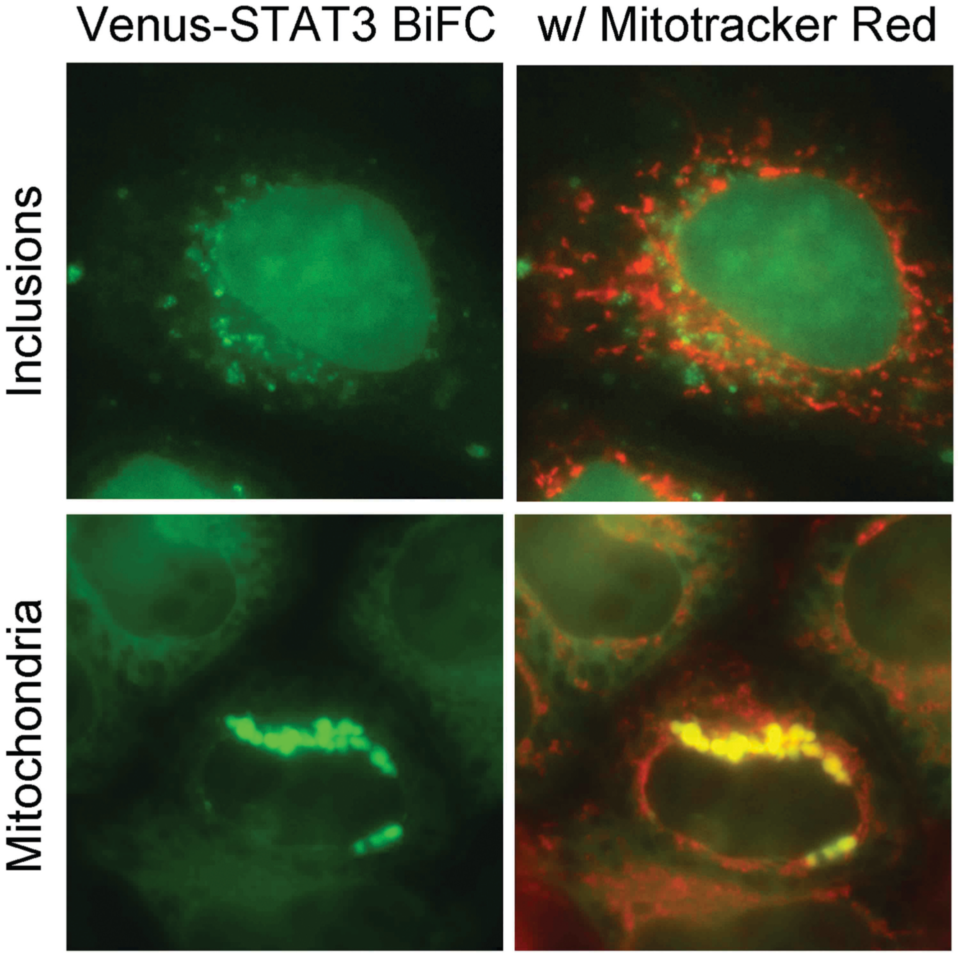
Venus-STAT3 BiFC
w/ Mitotracker Red
Inclusions
Mitochondria

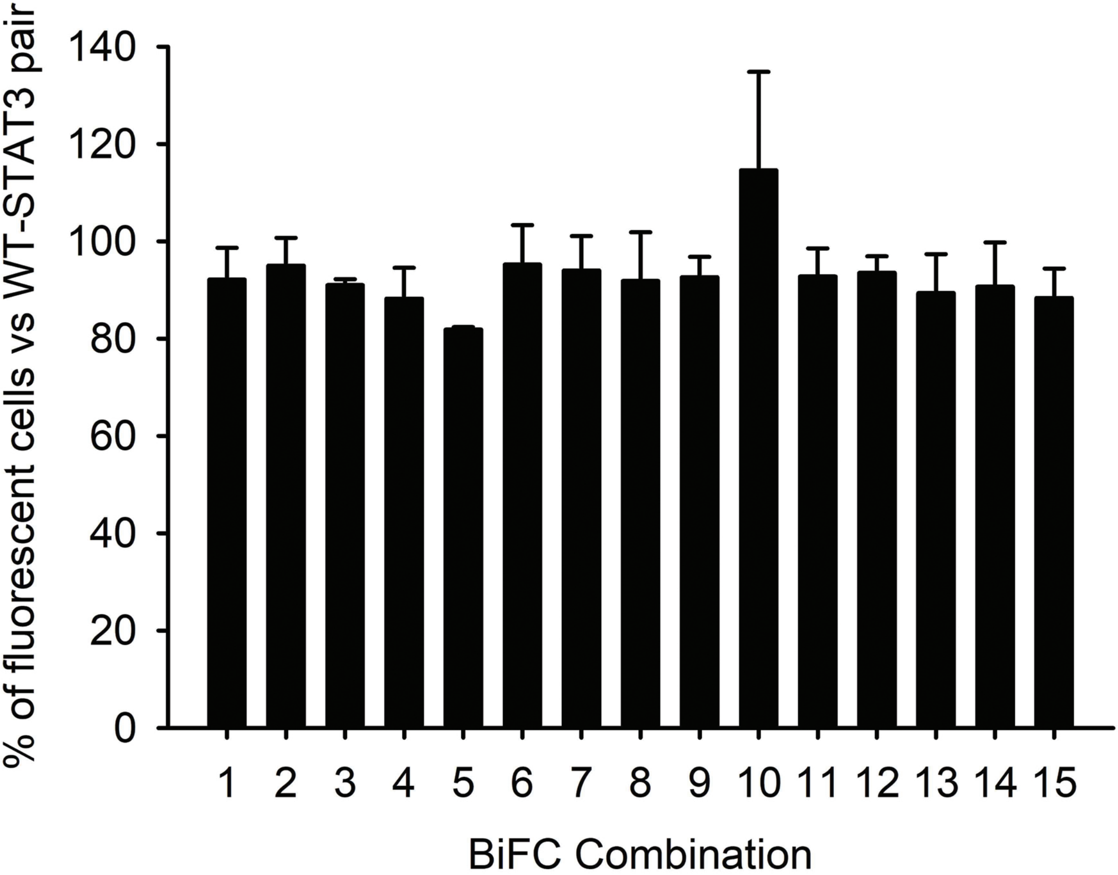


| | K49R | K140R | K685R | Y705F | S727A |
|---|---|---|---|---|---|
| K49R | 1 | | | | |
| K140R | 2 | 6 | | | |
| K685R | 3 | 7 | 10 | | |
| Y705F | 4 | 8 | 11 | 13 | |
| S727A | 5 | 9 | 12 | 14 | 15 |
